# Electrophysiological decoding captures the temporal trajectory of face categorization in infants

**DOI:** 10.1101/2024.10.07.617144

**Authors:** Roman Kessler, Michael A. Skeide

## Abstract

The adult human brain rapidly distinguishes between faces at around 170 milliseconds after stimulus onset. In the developing brain, however, the time course of face discrimination is poorly understood. To shed light on this issue, we presented human and nonhuman primate faces to five to thirteen-month-old infants in an event-related electroencephalography experiment. Using time-resolved decoding based on logistic regression we detected above-chance discrimination of human faces from nonhuman faces in a time window starting at around 200 milliseconds, originating from occipito-temporal electrodes. There was no evidence, however, for above-chance discrimination of individual human or individual nonhuman faces. Moreover, using neural network-based decoding, we delivered the proof of principle that face categorization but not individuation can be detected even at the level of single participants. These results indicate that rapid face categorization emerges already in preverbal infants.

## Introduction

Spatially and temporally selective responses to faces compared to other visual objects are long known in the adult human brain^1–3^. Moreover, perceiving different face categories, such as human versus nonhuman primate faces, is known to induce dissociable brain responses in the electroencephalogram. Specifically, the N170, an event-related potential peaking at around 170 milliseconds (ms) over occipito-temporal sources, exhibits shorter peak latencies for human faces compared to nonhuman primate faces^4,5^.

Spatially and temporally selective responses to faces in contrast to other object categories have also been observed in the developing infant brain^6–8^. In contrast to adults, however, there is no conclusive evidence for the existence of a face-category-selective N170 response in infants^9^. This being said, it has been shown that nine-month-old infants detect human or nonhuman primate faces presented as deviant stimuli within a series of standard faces from the other species category^10^. Furthermore, one study suggests that separating human faces from cat faces elicits an N290 response peaking at around 290 ms over bilateral occipital sources in infants at seven months of age^11^. This event-related potential has also been identified by a number of other infant studies focusing on object categorization by comparing faces with non-facial objects^12–14^. The time course of perceiving different face categories, however, remains poorly understood in the developing brain.

To resolve this challenge, we developed an event-related stimulus presentation design in which 38 five-to thirteen-month-old infants were exposed to human and nonhuman primate faces during electroencephalographic recording. The purpose of this design was to investigate how the infant brain discriminates between face categories (human primate vs. nonhuman primate) as well as individual faces within the same category (human 1 vs. human 2 and monkey 1 vs. monkey 2). Face discrimination was identified in the electrophysiological time course using time-resolved decoding based on logistic regression. The rationale for choosing time-resolved decoding instead of event-related potentials was to create a feature space that includes the electrophysiological information of all recording electrodes instead of modeling each electrode as a single isolated signal source^15,16^. In addition, for single-participant data we applied a convolutional neural network decoder known to outperform other decoding frameworks in within-participant analyses^17^.

Based on two previous event-related potential studies, we hypothesized to detect discrimination of face categories around 260 to 340 ms in occipito-temporal electrodes^9,11^. The discrimination of individual faces was expected in a slightly later time window than the discrimination of face categories^18,19^.

Contrary to our prediction, we found above-chance decoding of human versus nonhuman primate faces as early as 200 ms in occipito-temporal electrodes. At the same time, decoding of individual human and nonhuman primate faces remained at chance level. Neural network-based decoding of single-participant data also revealed significant face categorization in the majority of infants while there was no evidence for face individuation.

## Results

### Time-resolved decoding

At the group level, class labels were decoded for each time point separately from trial-wise whole-brain electrophysiological data using time-resolved logistic regression. Decoding was conducted for each participant individually, based on three comparisons: face categorization (human vs. monkey), human face individuation (human 1 vs. human 2), and monkey face individuation (monkey 1 vs. monkey 2). Across participants, human versus monkey faces were classified significantly above chance level for most time points between 200 and 770 ms (Fig. 1A, *p* < 0.05, one-sided, family-wise-error-corrected). No temporal cluster exceeded the significance threshold when decoding single individuals within a species (Fig. 1B,C).

**Fig. 1.**
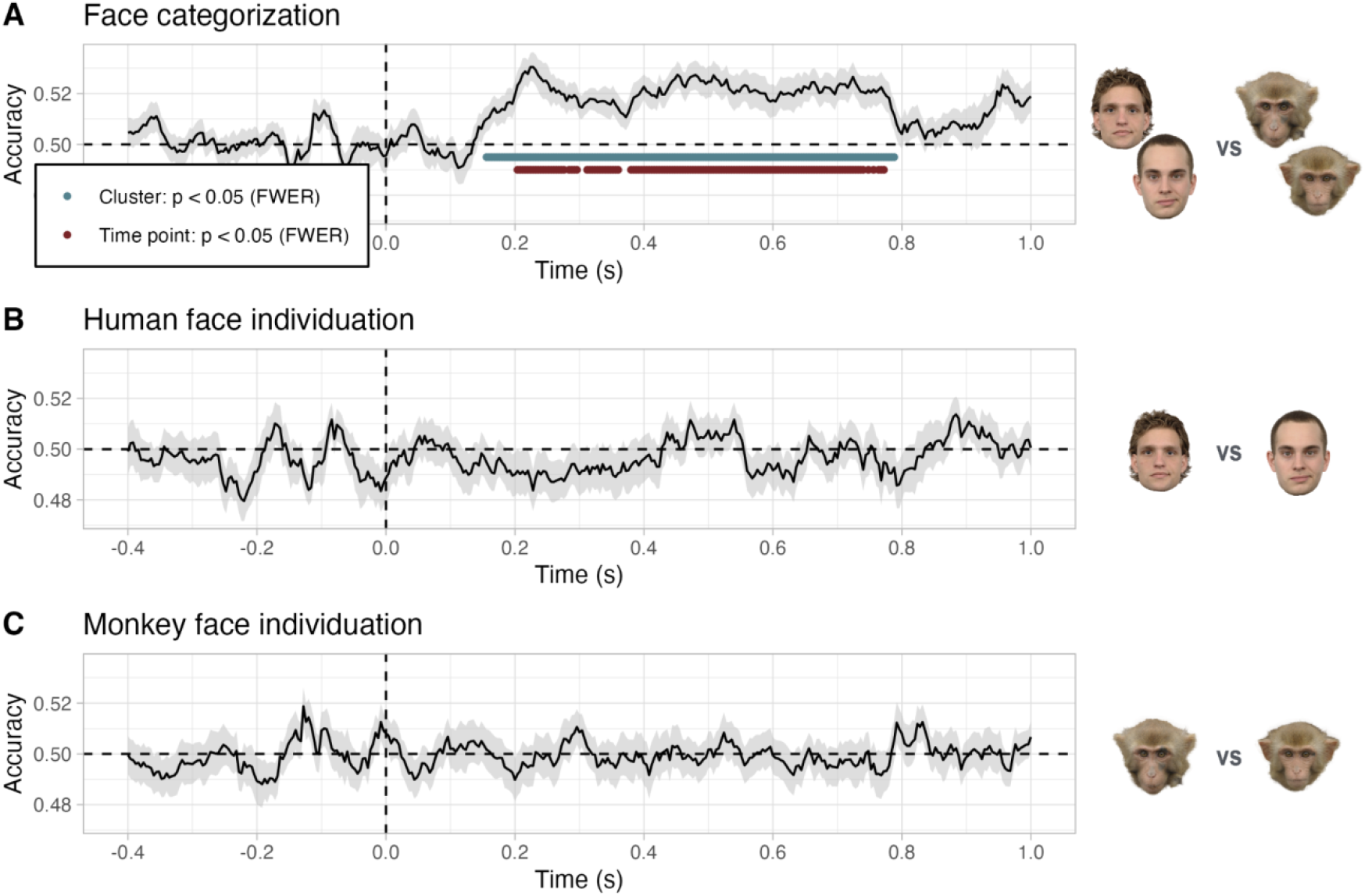
Time-resolved decoding accuracy at the group level. **A**, Face categorization. **B**, Human face individuation. **C**, Monkey face individuation. Cross-validated test accuracies (y-axes) are illustrated for each time point (x-axes), averaged across participants. Shaded gray areas depict the standard error of the mean. Dashed vertical lines depict stimulus onset. Dashed horizontal lines depict chance level. Blue-green dots illustrate significant temporal clusters based on a cluster mass test (*p* < 0.05, one-sided, family-wise-error-corrected). Dark red dots illustrate significant time points based on a cluster depth test (*p* < 0.05, one-sided, family-wise-error-corrected).

We also estimated the spatial electrophysiological response patterns over time across participants. Figure 2 illustrates the effect strengths for different time points and electrodes corresponding to face categorization (human vs. monkey comparison) shown in Figure 1A. Face categories were separable in bilateral occipital electrodes and in posterior temporal electrodes but with weaker activation and a slight right-hemispheric dominance (Fig. 2).

**Fig. 2.**
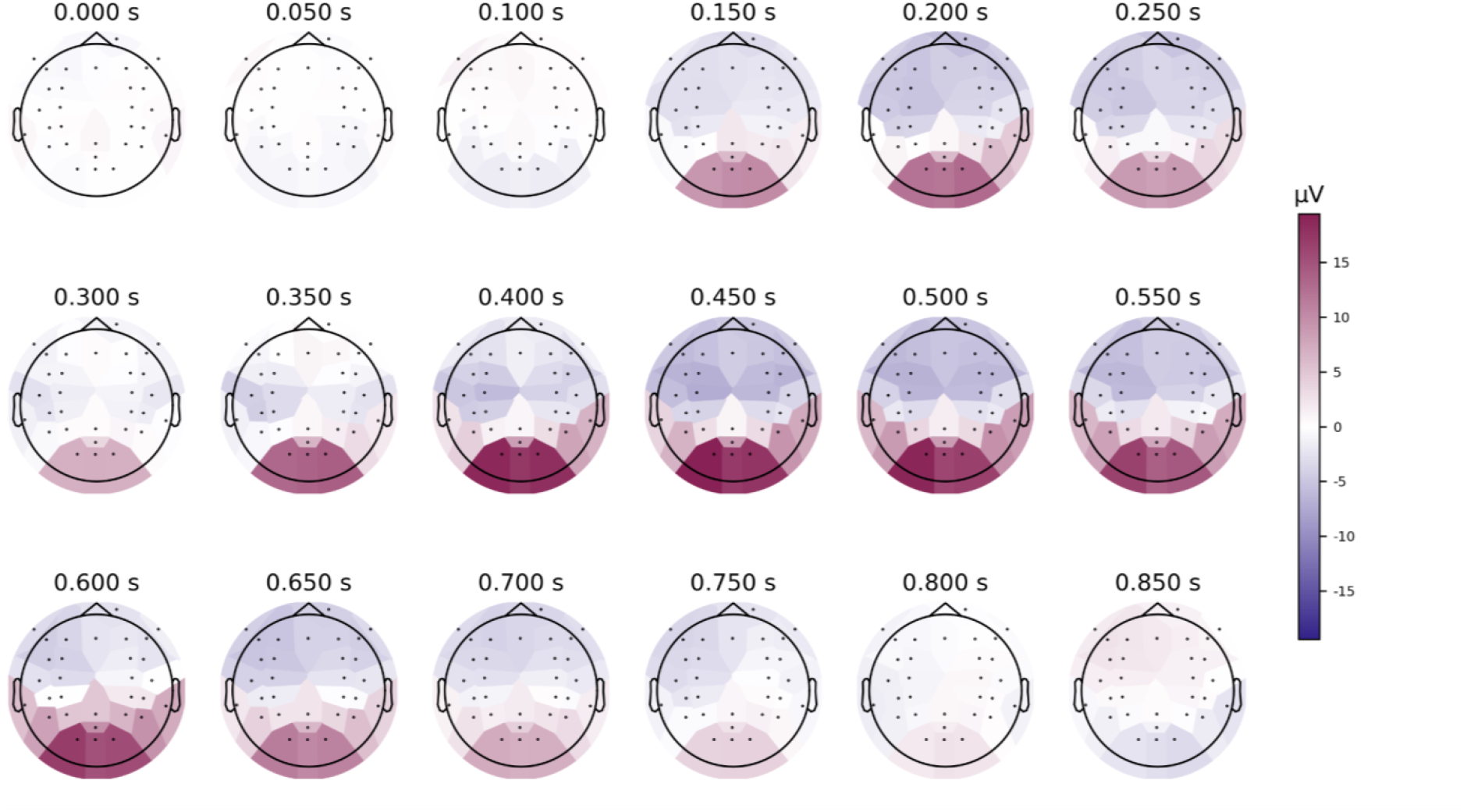
Feature importance maps for group-level time-resolved decoding of face categories. Spatial electrophysiological response patterns averaged across participants were calculated for several post-stimulus time points. Strongest feature importance for face categorization (human vs. monkey) was detected in bilateral occipital electrodes. Posterior temporal electrodes revealed weaker feature importance and a slight right-hemispheric dominance. µV denotes the effect strength in microvolts.

### Neural network-based decoding

At the level of single participants, we employed a convolutional neural network to decode class labels across time points from trial-wise whole-brain electrophysiological data, separately for each participant. Shuffled-label permutation tests revealed significant face categorization (human vs. monkey) in 21 out of 38 participants (*p* < 0.05, one-sided, false discovery rate-corrected) (Fig. 3A). Face individuation, however, remained at chance level in all participants (Fig. 3B,C). There was no evidence for a significant association between decoding accuracy and the number of trials available for each participant (Spearman’s rank correlations; face categorization *ρ* = 0.014, *p* = 0.93, human face individuation ρ = -0.168, *p* = 0.31, and monkey face individuation *ρ* = -0.004, *p* = 0.98). For the face categorization comparison, we evaluated the impact of single stimuli (i.e., human 1 & 2, and monkey 1 & 2) on decoding accuracy across participants. That is, we tested whether the network learned trials of particular stimuli better than trials of others. There was no evidence for performance differences between individual stimuli (paired *T*-tests, two-sided, all *p* > 0.2, false discovery rate-corrected).

**Fig. 3.**
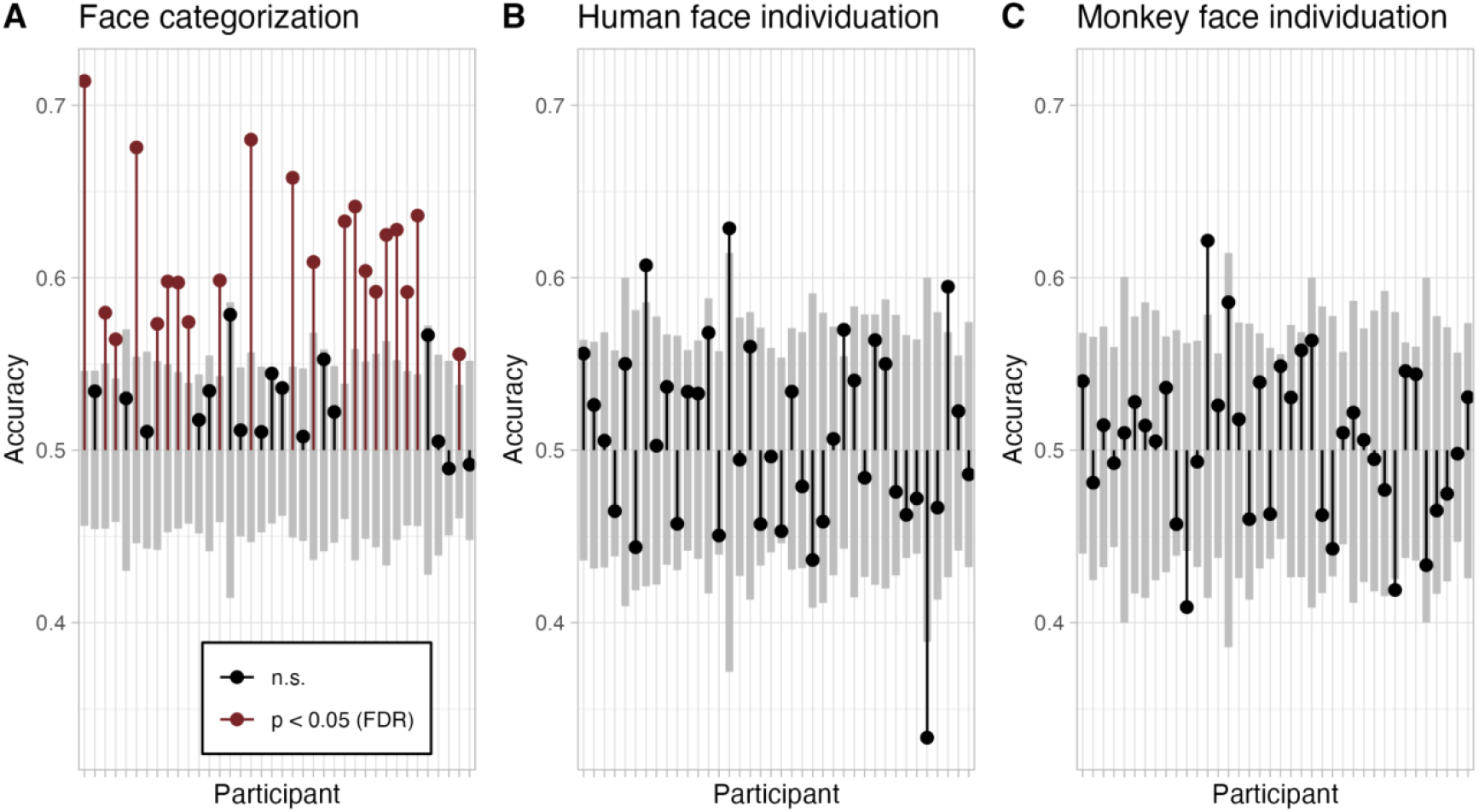
Neural network-based decoding accuracy in single participants. **A**, Face categorization. **B**, Human face individuation. **C**, Monkey face individuation. Cross-validated test accuracies (y-axes) are illustrated for each participant (x-axes) with respect to chance level (0.5). Red lines and dots depict participants in which decoding accuracy was significantly above chance (*p* < 0.05, one-sided, false discovery rate-corrected). Gray bars indicate the distribution of surrogate decoding accuracies (0.05 to 0.95 quantile). FDR: false discovery rate, n.s.: not significant.

Population prevalence of decoding above chance level was estimated using uncorrected *p*-values of each single participant and comparison. The posterior density of the population prevalence revealed a maximum a posteriori (MAP) estimate of 0.56, with a 95% highest density interval (HDI) ranging from 0.39 to 0.71 for face categorization (human vs. monkey). For human face individuation, MAP was estimated to be 0.06 (HDI: 0.00–0.18), while for monkey face individuation, MAP was estimated to be 0.00 (HDI: 0.00–0.09). Accordingly, decoding face categorization above chance level was feasible for an estimated 56% of the population, while within the current experimental framework, decoding human and monkey face individuation was only feasible for 6% and 0% of the population, respectively.

## Discussion

Time-resolved logistic regression-based decoding indicated that the infant brain discriminates between face categories at around 200 to 770 ms. In contrast, individual faces within the same category were not separable above chance level. Convolutional neural network-based decoding revealed face categorization but not face individuation in the majority of infants.

Going beyond event-related potentials, our time-resolved decoding approach provides evidence that face categorization in preverbal infants starts already around 200 ms. This early time window has a later onset than the adult N170 but precedes the N290 found at a preverbal infant age^4,5,9,11^. Differences between the present study and the previous studies in adults might be explained by slower encoding of face configuration information due to only gradually emerging fast electrophysiological activity^20–22^. The detection of an earlier time window compared to event-related potentials in infants could be attributed to the considerably larger number of trials collected here. Moreover, decoding makes use of features from all available electrodes, which can be expected to increase sensitivity^23^. That being said, contributions of several other factors such as the particular physiological age of the participants, the specific stimulus material selected or preprocessing parameters chosen are also possible (Kessler R., Enge A., Skeide M.A., in preparation). Accordingly, follow-up work is needed to replicate these results and to determine whether the effect generalizes to stimuli including other gender, ethnicity, or species.

Face category discrimination extended to time points as late as 770 ms which could hypothetically be related to the semantic differentiation of face categories after accessing semantic knowledge stored in long-term memory. Enabling semantic processing while continuously processing incoming stimuli also requires sustained attention. Both functions have been consistently ascribed to the infant Nc, an event-related potential peaking between around 300 and 800 ms over occipito-parietal and frontal sources^14,24–26^.

In the present dataset, several occipito-temporal electrodes contribute to the decoding accuracy while posterior occipital electrodes show particularly high feature importance, indicating a substantial contribution of low-level visual processing components. Additional experiments manipulating high-level semantic information while controlling low-level visual stimulus features are needed to narrow down the functions of this extended late stage of face categorization. It is not clear, however, whether such manipulation could be operationalized in an experiment for preverbal infants given that they are not able to solve behavioral semantic tasks. Moreover, manipulating images to match low-level features, such as spatial frequency or brightness, introduces changes with respect to other low-level features that could still contribute to classification performance. At the same time, it is not clear whether it would be feasible to match all low-level features while preserving ecological validity by keeping all facial features.

Employing neural network-based decoding we demonstrated the feasibility of detecting electrophysiological face category discrimination in single infants. To this end, we collected a comparably large number of up to 636 trials after quality control within relatively short recording sessions of less than thirty minutes in total on average. Our approach thus holds promise to facilitate the translation of electroencephalographic recordings into clinical settings. For example, the current results could stimulate further research on the early detection of neurodevelopmental disorders associated with impaired face recognition, in particular autism spectrum disorders^27–29^.

## Methods

### Participants

44 participants were invited for one or two sessions. Six participants did not achieve the predefined number of trials and were therefore excluded from the analysis (>20 trials per category and session or >35 trials per category in one session). Valid trials were determined by visual inspection of videos showing whether the infant looked at the stimuli or not. The remaining 38 participants (19 female, 19 male) had a mean age of around 8.2 months (range: 5–13 months). The total number of sessions was 71 and the mean interval between two sessions was 3.4 months. All caregivers gave written informed consent to participate. The study was approved by the Ethics Committee of the Medical Faculty of the University of Leipzig, Germany (149/22-ek). Data acquisition took place at the Max Planck Institute for Human Cognitive and Brain Sciences in Leipzig, Germany, between May 2023 and May 2024.

### Stimulus selection

We presented faces of two human individuals and two monkey individuals, including each individual with two different images. The final set of stimuli was limited to eight images to maximize the number of trials presented given the time constraints of an infant experiment while still being able to generalize beyond the particular features of a single stimulus. Human faces were chosen from the Radboud Faces Database^30^. Only white, male individuals were chosen. To achieve slight variability in the stimulation, but avoid strong emotional facial expressions, we initially chose three candidate individuals for which the neutral and the contemptuous facial expression looked very similar, i.e., the contemptuous facial expression looked relatively neutral. The faces were cropped by removing everything below the chin, but keeping characteristics such as hair and ears. Rhesus macaque monkey individuals were selected from the PrimFace face database of nonhuman primates (http://visiome.neuroinf.jp/primface). The monkeys initially chosen looked frontally into the camera. Some monkeys were excluded due to very different appearance, such as having a very different skin or fur color. Only monkey individuals with at least two different face images available were initially selected, resulting in six candidate individuals. The images were rotated so that the eyes were aligned horizontally.

We used AlexNet, a convolutional neural network^31^, to estimate the similarity between low-level visual features within and between individuals of a species. AlexNet was chosen as its layers are supposed to follow roughly a brain-like processing hierarchy^32,33^. We extracted and concatenated the AlexNet activations of the first five convolutional layers. Linear correlation coefficients were calculated between the activations of each stimulus candidate pair. First, we aligned within-individual similarity by choosing images with comparable correlations within each category (human or monkey). We only continued with individuals showing a comparable similarity between images. Then we aligned between-individual similarity within a species, by choosing two individuals of each species with comparable similarity estimates. This procedure mitigates the possibility that differences in decoding accuracy for the two face individuation comparisons (see below) are caused by a larger low-level visual feature difference between individuals within one category (human faces or monkey faces). Four individuals (two humans, two monkeys) were finally presented using two different images for each individual.

### Experimental design

Each image was presented repeatedly in a pseudo-randomized and event-related fashion. Faces were surrounded by a gray background and located centrally on the screen, occupying a visual angle of approximately 10°. At each presentation event, the stimulus randomly varied in size (80-100% of the dedicated visual angle) and orientation (±5% relative to origin). Each stimulus was presented for 750 ms, with a variable inter-stimulus-interval of 600 to 900 ms, drawn from a uniform distribution. Colorful videos were displayed occasionally whenever the participant’s focus on the screen faded or motivation decreased. Similarly, breaks were introduced by the experimenter when the participant’s attention started to go down. The experiment ended as soon as the participant started to become fuzzy, expressed discomfort, or after a maximum of 720 stimulus presentations (60 repetitions of each image), corresponding to 18 minutes of continuous stimulation.

### Data acquisition

The participants were standing or sitting on their caretaker’s lap in front of the screen. Both were continuously surveilled by a camera. The light in the room was dimmed and the room temperature was controlled to avoid sweating under the electroencephalogram (EEG) cap. Active electrodes (Brain Products, active electrodes) were positioned using a custom montage with 34 electrodes (31 EEG: Fz, F3, F7, F9, FC5, POz, C3, T7, TP9, CP5, P7, P3, Pz, O1, O2, P8, P4, CP6, TP10, T8, C4, Oz, FC6, F10, F8, F4, Fp2, FC4, CP4, FC3, CP3; one EOG; reference FCz; and ground Fp1). The EOG electrode was positioned on the right cheek of the participant. EEG was recorded using BrainVision Recorder (v. 1.22.0001, Brain Products) at a sampling rate of 1000 Hz with a low-pass filter of 70 Hz. Caps (Easycap) were pre-gelled before being placed on the participant’s head to minimize preparation time. Additional gel (custom-made: hydroxyethyl cellulose, propylene glycol, propylene glycol, sodium chloride, water) was applied to the electrodes until impedance was below 50 kΩ, but not for longer than 2-3 minutes, depending on the participant’s compliance.

### Data preprocessing

Data was mainly processed using *Bash* and *Python* (v. 3.11.6) on a high-performance computing cluster of the Max Planck Computing and Data Facility (Garching, Germany). Preprocessing of electrophysiological data was performed using *MNE* (v. 1.5.1)^34^. Two raters inspected the video data, and each trial was included if one rater was certain that the participant attended the screen during the trial. The trials rated as non-attentive by both raters were removed from the data. Only participants completing at least 20 trials from each condition in each experimental session, or 35 trials in a single session were included. The raw data was resampled to 250 Hz. Two artificial EOG channels were created, (1) by subtracting the electrode voltages of the physical EOG channel (below the right eye) and channel Fp1 (above the right eye), and (2) by subtracting F9 and F10. The physical EOG channel was then removed from the data.

The raw time series was high pass filtered at 0.5 Hz and low pass filtered at 15 Hz using a linear finite impulse response filter. An independent component analysis based on the picard method was conducted on a copy of the data filtered at 1 Hz, estimating 20 components in up to 500 iterations. Each component was correlated with the two artificial EOG channels. A component was classified as an ocular artifact if the correlation surpassed a threshold estimated via adaptive z-scoring. The solution was then applied to the original data to remove those artifactual components. A robust average was chosen as a common reference as implemented in *pyprep*^*35*^. Epochs were created using a pre-stimulus baseline window of 200 ms, and spanning a time window until 1000 ms after stimulus onset. Artifact correction was conducted using the *autoreject* package^36,37^. To not further reduce the number of trials, *autoreject* hyperparameters were set to avoid rejection of entire trials, but instead to interpolate channels in noisy trials (Kessler R., Enge A., Skeide M.A., in preparation). We calculated the amount of trials available for each of the four individual faces. Surplus trials were then randomly removed from the majority classes to achieve an equal number of trials per class. The preprocessed trials of both sessions were finally concatenated after trial balancing to avoid decoding the session instead of the class.

### Decoding analyses

We conducted both time-resolved and neural network-based decoding to predict stimulus class from the electrophysiological data. Decoding was performed based on three comparisons: face categorization (human vs. monkey), human face individuation (human 1 vs. human 2), and monkey face individuation (monkey 1 vs. monkey 2). Classifiers were trained using the data of each participant individually.

#### Time-resolved decoding

For time-resolved decoding as implemented in *MNE*, we standardized voltage values separately for each time point and channel across epochs. Next, we trained separate classifiers for each stimulus-locked time point. Then we employed a logistic regression classifier using liblinear solver and L2 penalty^38^. Subsequently, we conducted a stratified 10-fold cross-validation. The test accuracies were finally averaged across folds but separately for each time point, resulting in one decoding time series per participant and per comparison. We also quantified the feature importance for the face categorization comparison by multiplying classifier weights with the data covariance to retrieve spatial activation patterns^16,39^. For each time point, we then averaged the activation patterns across participants.

#### Neural network-based decoding

Full-trial-wise decoding was performed using *EEGNet* v4^17^ as implemented in the *braindecode* toolbox^40^. EEGNet is a convolutional neural network, a nonlinear method that extracts spatial and temporal features from the EEG signal to perform classification. In this procedure, entire trials of both classes entered the network and underwent several convolutions to finally predict the stimulus class. For signal rescaling, an exponential moving standardization was applied to each epoch beforehand. Each network was trained for 200 cycles with default parameters and a batch size of 16. We used a stratified 10-fold cross-validation and averaged the test accuracies across folds, resulting in one accuracy estimate per participant and comparison.

### Statistical analyses

Statistical analyses were conducted using *targets*^*41*^ in *R* (v. 4.2.2, https://www.R-project.org/) on Mac.

#### Time-resolved group analysis

We employed a permutation cluster mass test to delineate significant temporal clusters that were decodable above chance across participants^42,43^. In addition, we used a permutation cluster depth test to delineate significant time points^44^. The cluster mass test corrects the family-wise-error-rate at the cluster level, whereas the cluster depth test corrects the family-wise-error-rate at the level of individual timepoints. We used a cluster-forming threshold of 0.01 (1% above chance), and 10,000 permutations. We used permutation-based tests as they are independent of assumptions about data distributions.

#### Single-participant statistics on neural network test accuracy

To calculate statistics based on each participant’s individual test accuracy per comparison, we conducted shuffled-label permutation testing^45^. Over 1000 iterations, we shuffled the class affiliation of trials of the train and test sets and used them to estimate surrogate accuracies. A participant’s test accuracy of the correctly labeled data was then related to the surrogate test accuracies of the Monte Carlo estimation of the permutation distribution to compute a *p*-value. We corrected for the false discovery rate across participants per comparison using the Benjamini-Hochberg^46^ procedure for visualization. Uncorrected *p*-values were used for estimating the population prevalence.

#### Population prevalence

We estimated the population prevalence of above-chance decoding with neural networks using Bayesian inference on population prevalence^47,48^. Bayesian prevalence estimates how many percent of participants sampled in the future will exhibit a significant effect at a threshold of *p* < 0.05 and a corresponding uncertainty estimate. For this purpose, we set an uncorrected alpha level of 0.05 on the accuracies of each participant obtained from neural network-based decoding, and derived a maximum a posteriori estimate and a 95% highest posterior density interval under a uniform prior for each comparison.

## Data availability

The data collected for this study is available through a public link on Zenodo (https://doi.org/10.5281/zenodo.13881207).

## Code availability

All code used for data analysis is available from GitHub (https://github.com/SkeideLab/PRAWN).

## Acknowledgements

We thank our lab manager Micha Vollmann for recruiting participants, coordinating logistics, handling forms, and organizing the measurements. This work was supported by the German Research Foundation (DFG Heisenberg Program Grant 433758790 awarded to M.A.S.) and the Jacobs Foundation (Research Fellowship awarded to M.A.S.). The funders had no role in study design, data collection and analysis, decision to publish or preparation of the manuscript. The rhesus monkey face images used in this study are provided by the PrimFace database: http://visiome.neuroinf.jp/primface, funded by a Grant-in-Aid for Scientific research on Innovative Areas, “Face Perception and Recognition” from the Japanese Ministry of Education, Culture, Sports, Science, and Technology (MEXT).

## Author contributions

M.A.S. conceived the study. R.K. performed the experiments and analyzed the data. R.K. visualized the results with feedback from M.A.S. R.K. and M.A.S. wrote the manuscript. M.A.S. supervised the study.

## Ethics declarations

The authors declare no competing interests.

