## Supplementary Material for "Electrophysiological decoding captures the temporal trajectory of face categorization in infants"

**Table S1 | Demographics and trials counts per participant.** Number of trials in each of the four stimulus conditions (human 1, human 2, monkey 1, and monkey 2) after quality control. Age refers either to the age in months at the experimental session, if only one session was included, or to the average age across sessions, if two sessions were included. Sex: m=male, f=female.

| Participant | Number of trials per condition | Age (months) | Sex |
|---|---|---|---|
| **sub-001** | 126 | 13 | m |
| **sub-002** | 108 | 8 | f |
| **sub-003** | 105 | 9 | f |
| **sub-004** | 135 | 9 | m |
| **sub-005** | 51 | 9 | m |
| **sub-006** | 82 | 7.5 | m |
| **sub-007** | 71 | 13 | m |
| **sub-008** | 84 | 7.5 | m |
| **sub-009** | 105 | 8.5 | f |
| **sub-010** | 113 | 7.5 | f |
| **sub-012** | 144 | 9 | f |
| **sub-013** | 115 | 9.5 | f |
| **sub-014** | 73 | 9 | f |
| **sub-015** | 139 | 9 | f |
| **sub-016** | 38 | 8 | m |
| **sub-017** | 98 | 7.5 | f |
| **sub-019** | 76 | 9 | f |

| | | | |
|---|---|---|---|
| **sub-020** | 109 | 8.5 | m |
| **sub-021** | 135 | 8.5 | f |
| **sub-022** | 150 | 7.5 | f |
| **sub-023** | 98 | 7 | m |
| **sub-024** | 96 | 8 | m |
| **sub-025** | 56 | 6 | m |
| **sub-026** | 76 | 7.5 | m |
| **sub-028** | 90 | 7 | m |
| **sub-030** | 159 | 8 | m |
| **sub-032** | 65 | 8.5 | m |
| **sub-033** | 86 | 8 | f |
| **sub-034** | 75 | 8.5 | f |
| **sub-035** | 66 | 8.5 | m |
| **sub-036** | 93 | 8 | f |
| **sub-038** | 121 | 8.5 | f |
| **sub-039** | 125 | 7 | m |
| **sub-040** | 48 | 5 | f |
| **sub-041** | 76 | 8 | m |
| **sub-042** | 99 | 8 | f |
| **sub-043** | 156 | 8 | f |
| **sub-044** | 93 | 8.5 | m |
